# Memory-paced tapping to auditory rhythms: Effects of rate, speech and motor-engagement

**DOI:** 10.1101/2021.07.13.452153

**Authors:** Anat Kliger Amrani, Elana Zion Golumbic

**Author notes:** <u>Corresponding Author:</u> Elana Zion Golumbic, The Gonda Center for Multidisciplinary Brain Research, Bar Ilan University, Ramat Gan, Israel.

## Abstract

Humans have a near-automatic tendency to entrain their motor actions to rhythms in the environment. Entrainment is hypothesized to play an important role in processing naturalistic stimuli, such as speech and music, which have intrinsically rhythmic properties. Here we studied two facets of entraining one’s rhythmic motor actions to an external stimulus: (1) synchronized finger tapping to auditory rhythmic stimuli, and (2) memory-paced reproduction of a previously heard rhythm. Using modifications of the Synchronization-Continuation tapping paradigm, we studied how these two rhythmic behaviours were affected by different stimulus and task features. We tested synchronization and memory-paced tapping for a broad range of rates, from sub-second to supra-second, both for isochronous tone-sequences and for rhythmic speech stimuli (counting from one to ten), which are more ecological yet less strictly isochronous. We also asked what role motor engagement plays in forming a stable internal representation for rhythms and guiding memory-paced tapping.

Results show that individuals can flexibly synchronize their motor actions to a very broad range of rhythms. However, this flexibility does not extend to memory-paced tapping, which is accurate only in a narrower range of rates, around ^~^1.5Hz. This pattern suggests that intrinsic rhythmic-defaults in the auditory/motor systems influence the internal representation of rhythms, in the absence of an external pace-maker. Interestingly, memory-paced tapping for speech rhythms and simple tones shared similar ‘optimal rates’, although with reduced accuracy, suggesting that internal constraints on rhythmic entrainment may generalize to more ecological stimuli. Last, active synchronization led to more accurate memory-paced tapping vs. passive listening, which emphasizes the importance of action-perception interactions in forming stable entrainment to external rhythms.

## 1. Introduction

Many natural stimuli, such as music, speech and biological motion, have intrinsically rhythmic properties. It has long been hypothesized that effective processing of rhythmic stimuli involves synchronizing one’s own internal rhythms and sometimes generating motor rhythms to match those present in the environment (‘the entrainment hypothesis’; Lakatos et al. 2008; Haegens and Zion Golumbic 2018; Rimmele et al. 2018; Jones 2019). A prevalent paradigm often used to study the interaction between sensory perception and motor production of rhythms is the Synchronization-Continuation tapping paradigm (Stevens 1886). The paradigm consists of two phases: a ***Synchronization*** phase, where participants are required to tap in synchrony with a sequence of rhythmic tones, followed by a ***Continuation*** phase where participants are asked to continue tapping at the same pace after the rhythmic stimulation has ceased (Dunlap 1910; Bartlett and Bartlett 1959; Wing and Kristofferson 1973; Flach 2005). This elegant paradigm has allowed researchers to study the behavioural and neural components of the action-perception loop that support human rhythmic behaviour (Serrien 2008; Repp and Steinman 2010; Merchant et al. 2011; Donnet et al. 2014; Bravi et al. 2017).

The two phases of the paradigm – synchronization and continuation – tap into different aspects of rhythmic behaviour. Synchronization relies critically on precise coordination and entrainment between the auditory and motor systems (Kornysheva and Schubotz 2011; Schwartze et al. 2011, 2016; Repp and Moseley 2012; Repp and Su 2013; Tierney and Kraus 2013), and inaccurate synchronization is associated with deficits in both low-level auditory encoding and in perception-action coupling (Phillips-Silver et al. 2011; Sowiński and Dalla Bella 2013; Tierney and Kraus 2013; Palmer et al. 2014; Nozaradan et al. 2016; Schwartze et al. 2016). Conversely, continuation-tapping is performed in the absence of any sensory input and therefore relies on an internal representation of the rhythm, established during the synchronization phase (Ivry and Richardson 2002). Accordingly, continuation-tapping, which is also referred to as ‘memory-paced’ tapping, relies on neural mechanisms that are strongly linked to temporal working memory and the internal representation of time, such as the basal ganglia and SMA (Nenadic et al. 2003; Lewis et al. 2004; Lustig et al. 2005; McNab and Klingberg 2008; Wiener et al. 2010; Toyomura et al. 2012; Teki et al. 2017; Koshimori et al. 2019; Marvel et al. 2019; Schmidt et al. 2019). In other words, continuation tapping is driven by auditory imagery of a previously-experienced rhythm, rather than by the rhythmic stimulus. The distinction between stimulus-driven (synchronization) and memory-paced (continuation) tapping has been shown to be critical for understanding the nature of rhythmic deficits in several clinical conditions, such as Parkinson’s and ADHD as well as some learning disabilities (Thaut et al. 2001; Tierney and Kraus 2014; Woodruff Carr et al. 2014; Hove et al. 2017; Harrison et al. 2018; Jaeger et al. 2018; Kliger Amrani and Zion Golumbic 2020a).

In the current study we delve into the nature of the internal representations that drive rhythmic synchronization-continuation performance and ask three specific questions: (1) Is the internal memory representation that guides continuation-tapping perceptual in nature, or does its accurate formation depend on prior ***motor-engagement*** during synchronization? (2) Are accurate synchronization and memory-paced tapping affected by intrinsic rhythmic-defaults? (3) To what degree does rhythmic synchronization-continuation behaviour generalize beyond simple, precise rhythms to more ***ecological rhythmic stimuli***, such as speech?

### 1.1 Internal representation of rhythms

Studying the nature of the internal representations formed for perceptual rhythms has important implications for understanding the mechanisms underlying internal time-keeping and temporal prediction (Rohenkohl et al. 2012; Zion Golumbic et al. 2012; Cravo et al. 2013; Morillon and Baillet 2017; Haegens and Zion Golumbic 2018; Zalta et al. 2020). Two alternative hypotheses can be considered: The first is that the internal representations of previously-perceived rhythms that guide memory-paced tapping are sensory in nature, and are derived from encoding the temporal regularities of the auditory input. From a neural perspective, this hypothesis is in line with the notion of auditory neural entrainment, which suggests that neural oscillations in auditory cortex entrain to rhythms in the environment, and adjust their momentary frequency and phase so that high excitability phases coincide with predictable auditory events (Lakatos et al. 2005, 2013; Schroeder and Lakatos 2009; Zion Golumbic et al. 2012; Henry and Herrmann 2014; Breska and Deouell 2017; Haegens and Zion Golumbic 2018; Rimmele et al. 2018). However, another possibility is that a purely sensory-derived memory representation of rhythm is not sufficient for its accurate reproduction, and that engagement of the motor system is critical for generating a stable internal representation for rhythms. In other words, that accurate memory-paced tapping during the continuation phase is a product of cooperation between the auditory and motor systems during the synchronization phase. This perspective is motivated by the strong mutual influence observed between auditory perception and motor action, that manifests, for example, in perceptual enhancements brought about by movement (Thaut et al. 2001; Schroeder et al. 2010; Chemin et al. 2014; Morillon et al. 2014; Park et al. 2015; Rimmele et al. 2018; Gale et al. 2021; Reznik et al. 2021), as well as in the near automatic tendency to synchronize one’s body-movements to rhythmic sounds (Large 2008; Roerdink et al. 2011; Repp and Su 2013; Tranchant et al. 2016; Assaneo et al. 2019; Damm et al. 2020). Auditory-motor interactions are also hypothesized to involve neural oscillations as the backbone supporting communication and coupling between auditory and motor cortices and coordinating rhythmic behaviour (Arnal et al. 2015; Breska and Deouell 2017; Morillon and Baillet 2017; Morillon et al. 2019; Abbasi and Gross 2020).

Given these two alternatives, the first goal of this study was to weigh the relative importance of motor engagement for accurate memory-paced tapping. Specifically, if memory-paced tapping is just as good after listening passively to rhythms as it is after actively synchronizing to them, this would suggest that the internal representation for rhythms is primarily sensory-driven. However, if motor-synchronization improves subsequent memory-paced tapping, this would point to an important role of active motor engagement in establishing an internal representation for rhythms.

Another aspect to consider with regard to synchronization and memory-paced tapping is the effect of rate. Previous studies have found that there is a relatively narrow range of rates for which memory-paced tapping is optimal, roughly around ^~^0.6 sec SOA (1.6 Hz). This range corresponds to the rates of spontaneous motor rhythms and perceptual auditory preferences (McAuley et al. 2006; McAuley 2010; Roerdink et al. 2011; Large and Gray 2015a; Scheurich et al. 2018; Zamm et al. 2018; Kliger Amrani and Zion Golumbic 2020b, 2020a), leading to the suggestion that they reflect physiological rhythmic-defaults of the auditory and/or motor systems. Given these presumed built-in default rhythmic preferences, here we further tested whether the potential effect of motor engagement manifests similarly across a broad range of rates (from sub-second to supra-second) or whether it interacts with presumed default rhythmic preferences.

### 1.2 Representation of speech rhythms

Much of the growing interest in the behavioral and neural mechanisms of rhythmic perception stems from its potential ecological-relevance, since many naturalistic stimuli have rhythmic properties. One of the most commonly mentioned stimulus in this regard is natural speech, as it contains temporal regularities across several time-scales (Rosen and Fourcin 1986; Rosen 1992; Ghitza 2011; Loukina et al. 2011; Ding 2016; Assaneo et al. 2019; Morillon et al. 2019; Poeppel and Assaneo 2020). Indeed, several studies have demonstrated a strong link between rhythmic performance (and training) and speech-processing/language-development capabilities, leading to suggestions that speech-processing may be assisted by mechanisms involved in rhythm perception and temporal predictions (Thaut et al. 2001; Corriveau and Goswami 2009; Lim 2010; Woodruff Carr et al. 2014; Cortese et al. 2015; Schön and Tillmann 2015). However, despite this theoretical enthusiasm, the fact that speech-rhythms are not strictly isochronous raises questions regarding the generalizability of theoretical notions that are derived from responses to simple isochronous rhythms (e.g. tone sequences) to the more ‘relaxed’ and variable rhythms of natural speech (Haegens and Zion Golumbic 2018; Keitel et al. 2018; Kösem et al. 2018; Doelling and Assaneo 2021).

One of our goals was to contribute to this ongoing discussion regarding the transferability of insights from the processing of strictly isochronous auditory stimuli to the ability to detect and synchronize to rhythmic aspects of speech. To this end, we exchanged the simple-tone sequences used in the classic synchronization-continuation task with a simple rhythmic sequence of speech: counting from 1 to 10. We tested whether participants achieved similar, better, or worse tapping accuracy to these naturally-rhythmic speech stimuli relative to isochronous simple tones. Admittedly, this adaptation of the synchronization-continuation task to speech-derived rhythms is by no means representative of natural speech processing, nor is tapping to speech a particularly ecological task (Lidji et al. 2011; Villing et al. 2011). However, it provides a well-controlled framework for assessing whether individuals can synchronize to- and form robust internal representations for speech-derived rhythms, as they can for simple isochronous rhythms. Moreover, given the above-mentioned default rhythmic-preferences observed for spontaneous motor rhythms and isochronous tones (Roerdink et al. 2011; Large and Gray 2015a; Scheurich et al. 2018; Zamm et al. 2018; Kliger Amrani and Zion Golumbic 2020b, 2020a), this setup allows us to assess whether these defaults are present for speech-derived rhythms as well (Assaneo et al. 2021). If we find that synchronization and memory-paced tapping to simple and speech-derived rhythms share similar optimal rates, this would support the existence of a common rhythmic mechanism, such as default neural oscillations, underlying a variety of ecologically-relevant rhythmic behaviors.

To achieve the goals described above, participants performed the synchronization-continuation task under several conditions. To determine the importance of motor-engagement for forming internal representations of rhythms, we compared memory-paced tapping after active synchronization vs. after passive listening to a reference rhythm. To determine whether rhythmic performance is affected by the type of stimulus, we compared synchronization and memory-paced tapping to rhythmic sequences of simple tones and to rhythmic speech-sequences of counting. Last, both of these questions were tested on a broad range of rates, from sub-second to supra-second SOAs, which allowed us to explore the dynamic range of rhythmic capabilities and to gauge the effects of rhythmic default preferences.

## 2. Materials and Methods

### 2.1 Experimental design

#### Participants

The experiment included 17 participants (11 women, 6 men, age range 21-32, mean 25.2, all right handed). The experiment was approved by the Bar Ilan IRB and participants gave written informed consent prior to starting the experiment. None of the participants reported any neurological or psychiatric clinical condition.

#### Experimental Apparatus

Participants were seated comfortably in a sound attenuated booth, and heard sounds through headphones (Sennheiser HD 280 pro). The experiment was programmed and controlled using PsychoPy software (www.psychopy.org). Finger taps were recorded using a custom-made tapper based on an electro-optic sensor.

#### Stimuli

All auditory stimuli were prepared using the softwares Audacity and Matlab (Mathworks). Tones sequences consisted of 10 repetitions of pure tones (440Hz, 30ms with ±5ms ramp up/down) presented at 10 different rates (SOAs: 0.25, 0.35, 0.45, 0.55, 0.65, 0.8, 1, 1.4, 1.8 and 2.2 seconds). To create the counting stimuli, we recorded a female speaker counting in Hebrew from one to ten while listening and synchronizing her speech to a metronome (0.8 sec SOA; 75 bpm). Note that in Hebrew, all digits except for one are bi-syllabic words, however they vary in length and in the position of the stressed syllable (see Figure 4). To create rhythmic counting stimuli in a range of rates, the original recording was cut into 0.8sec-long intervals around each digit. These segments were then either stretched or shrunk using the Paulstrech function (Audacity software) to create intelligible versions of each digit for all SOAs. These segments were then concatenated back to form rhythmic counting streams from 1 to 10 at all rates (using Matlab; Figure 1A, right panel). The stimuli can be experienced and downloaded from The Open Science Framework (https://osf.io/nc67q).

**Figure 1.**
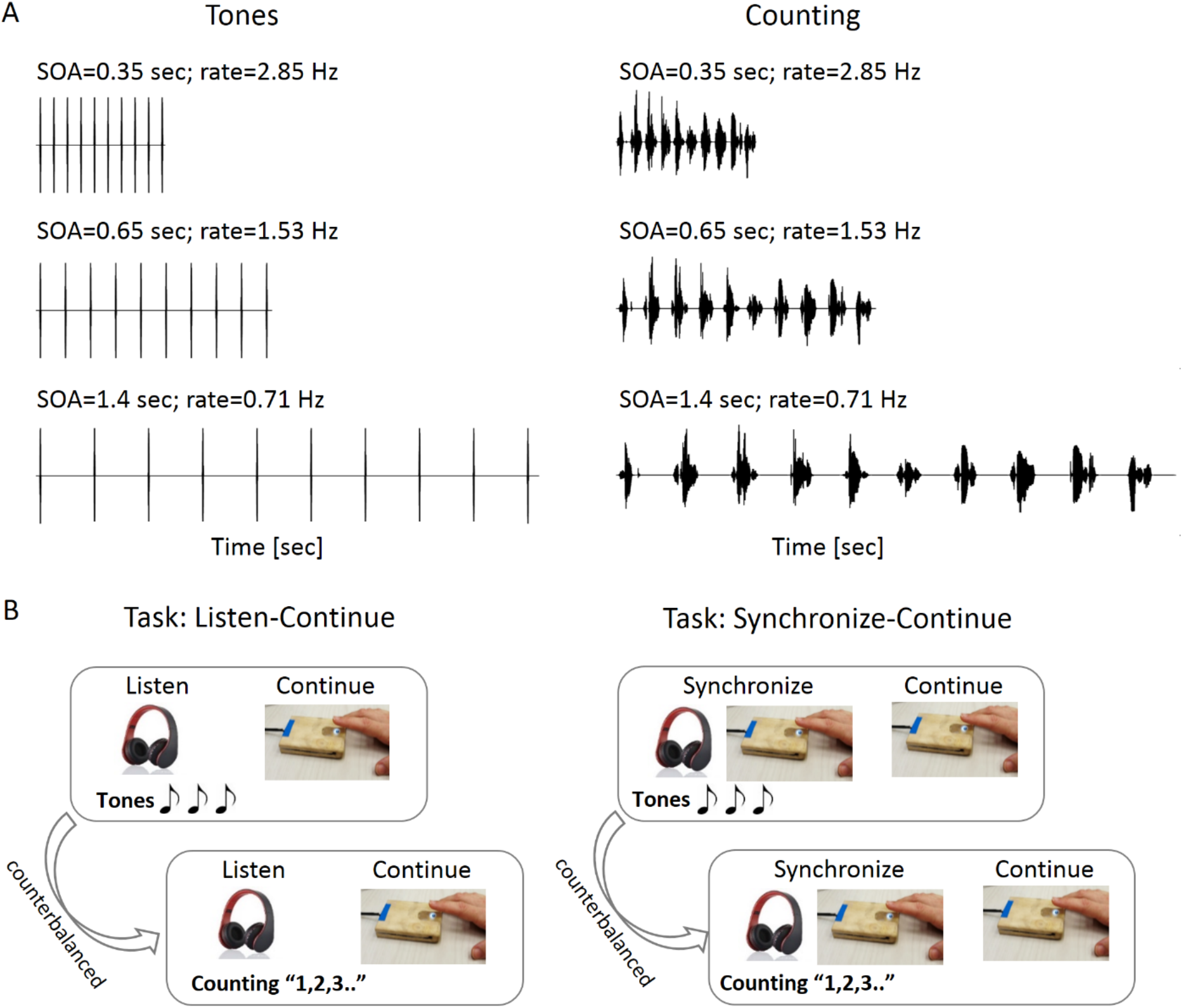
**A.** Examples of the Tone (left) and Counting (right) stimuli, at three different rates: SOA = 0.35 sec (top), SOA = 0.65 sec (middle) and SOA = 1.4 sec (bottom). **B.** Illustration of the experimental design. The Listen-Continue task always preceded the Synchronize-Continue task. Each task consisted of two Tone blocks and two Counting blocks, presented in counterbalanced order across participants.

#### Procedure

The experiment consisted of two consecutive tasks (Figure 1B):

##### Listen-Continue task

The Listen-Continue task is a passive-variation of the classic synchronization-continuation task. It consisted of a *Listen-phase* where participants heard a reference-rhythm, comprised either of ten simple tones or spoken digits (1-10). Following a 1.5 second-long pause, participants received visual instruction to start the *Continuation-phase* in which they were required to tap their finger exactly 10 times at same the rate they had just heard.

##### Synchronize-Continue task

This is the classic synchronization-continuation task. In the *Synchronization-phase* participants heard a reference-rhythm (ten simple tones or spoken digits) and were told to tap in synchrony with the auditory input they heard. Following a 1.5 second-long pause, participants received visual instruction to start the *Continuation-phase* which was identical to the Listen-Continue task, i.e. to tap their finger exactly 10 times at same the rate they had just heard.

Both the Listen-Continue and the Synchronize-Continue tasks were performed in blocks consisting of trials with different rates. The Listen-Continue task was always performed first, since we assumed that once an explicit synchronization task was performed this might contaminate subsequent Listen-Continue tasks. Within each of the tasks, Tones and Counting stimuli were presented in separate blocks (2 blocks per stimulus-type per task) and each block contained two repetitions of each of the ten rates used (random order). The order of tones vs. counting blocks within each task was counterbalanced across participants (Figure 1B). Participants did not receive feedback regarding the temporal accuracy of their continuation tapping.

### 2.2 Data Analysis

Two measures – precision error and tapping isochrony – were derived from the tapping times and Inter-Tap-Intervals (ITIs) measured in both the synchronization and continuation phase.

These measures capture complementary aspects of tapping accuracy, as described in our previous work (Kliger Amrani and Zion Golumbic 2020b, 2020a) (Figure 2).

**Figure 2.**
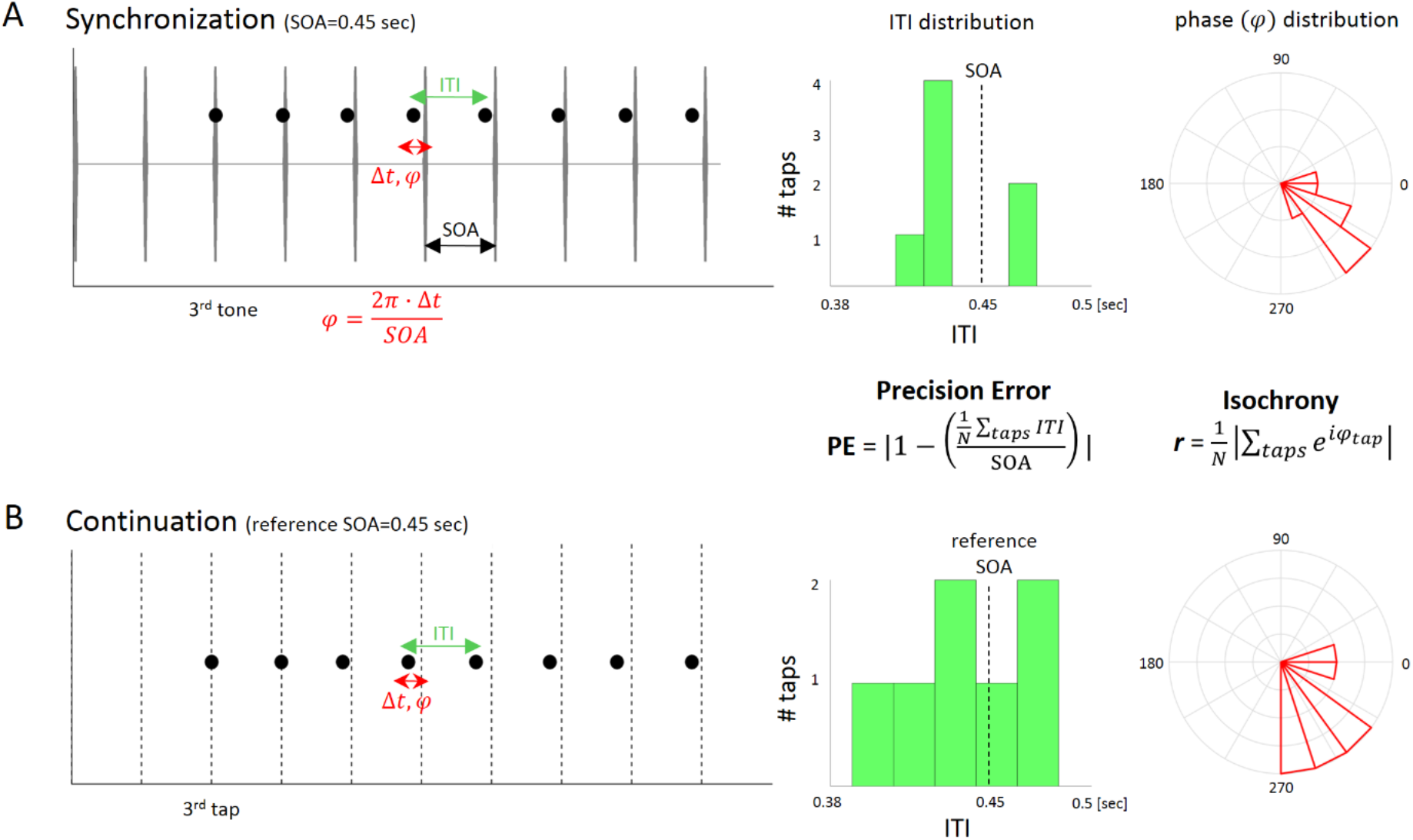
Illustration of the Precision Error and Isochrony measures. The example shows tapping data from a single trial in the Synchronization (A) and Continuation (B) phases, to a Tone stimulus (SOA = 0.45 sec). **(A) Synchronization phase**. Left: The rhythmic stimulus is shown in gray, and tap-times are indicated by black circles. The stimulus onset asynchrony (SOA; black arrow) is defined by the time between consecutive tone sounds. The inter-tap-interval (ITI; green arrow) is the time interval between each two consecutive taps, and was calculated for all tap-pairs (after removing the first two taps). ITIs may vary within a trial, as shown in the distribution of ITI values (middle). The **Precision Error** (PE) reflects the distance between the mean ITI in a given trial and the prescribed SOA of the stimulus. The phase of each tap (φ; red) is assessed by dividing the time-difference between each tap and the closest sound (Δt) by the interval length (SOA). The circular distribution of phase values (right) is used to calculate the resultant vector (r) which reflects the degree of tapping **isochrony**. **(B) Continuation phase.** Precision Error and Isochrony were also assessed for continuation tapping. ITIs and the resulting PE were assessed in the same manner as in the synchronization phase. Given the absence of an external reference rhythm, tap phases were assessed relative to an imaginary rhythm (dotted lines) at the rate of the reference rhythm that started at the participants’ 3^rd^ tap (first two taps were removed). Isochrony was calculated based on these phase values.

#### Precision error

captures the extent to which the average tapping rate deviates from the prescribed rate. Precision error (PE) is derived using the ratio between the mean ITI produced in 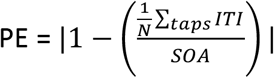, with lower values of precision error indicating a more precise replication of the required mean rate. As such, precision error can be considered a ‘global’ measure of tapping accuracy.

Complementing the precision error measure, the **Tapping Isochrony** measure captures the extent to which the intervals between taps are consistent throughout a trial. It is derived by quantifying the phase-shift between each individual tap and the prescribed rhythm, and calculating the consistency of phase-shifts across taps (Phase locking resultant; 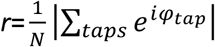, *circ_r.m* function, Matlab circular statistics toolbox). For synchronization tapping, the phase-shift is calculated relative to the start time of the acoustic stimulus. For continuation tapping, where there is no acoustic stimulus, we calculated the phase-shift of taps relative to an imaginary ‘reference rate’ at the prescribed rhythm (see Figure 2). Precision error and tapping isochrony were calculated separately for each tempo, condition and participate. To allow some time for establishing a stable rhythm, the first two taps in each trial were removed.

### 2.3 Statistical analysis

Since both the precision error and the isochrony measures are normalized, they are comparable across rates. Comparison of tapping accuracy during the synchronization phase was performed using a two-way Repeated Measures ANOVA, separately for the precision error and isochrony measures, with the factors: Stimulus (Tones/Counting) x Rate (10 rates). Note that this analysis was irrelevant for the Listen-Continue task, since it did not include synchronized tapping. For analysis of memory-paced tapping accuracy in the continuation phase, we performed a threeway Repeated Measures ANOVA on the precision error and isochrony measures. The three factors were: Task (Listen/Synchronize) x Stimulus (Tones/Counting) x Rate (10 rates). All statistical analyses were carried out using the JASP software (https://jasp-stats.org).

## 3. Results

### 3.1 Synchronization-Tapping

In line with previous studies, during the Synchronization phase tapping precision errors were extremely low and isochrony was high for most rates. There was a significant main effect of rate on both precision error [F(9,144)=4.38, p<0.001] and isochrony [F(9,144)=25.57, p<0.001]. This effect was driven by worse performance at the two fastest rates (Figure 3), however for all rates with SOAs slower than 0.5 sec synchronization precision errors and isochrony were steadily good. There was also a main effect of stimulus type on both precision error [F(1,16)=6.28, p=0.023] and isochrony [F(1,16)=29.99, p<0.001] indicating significantly better synchronized tapping when listening to Tones vs. Counting stimuli. The interaction between Stimulus Type x Rate was not significant for either measure [precision error: F(9,144)=0.61, p=0.79, isochrony: F(9,144)=0.52, p=0.86].

**Figure 3.**
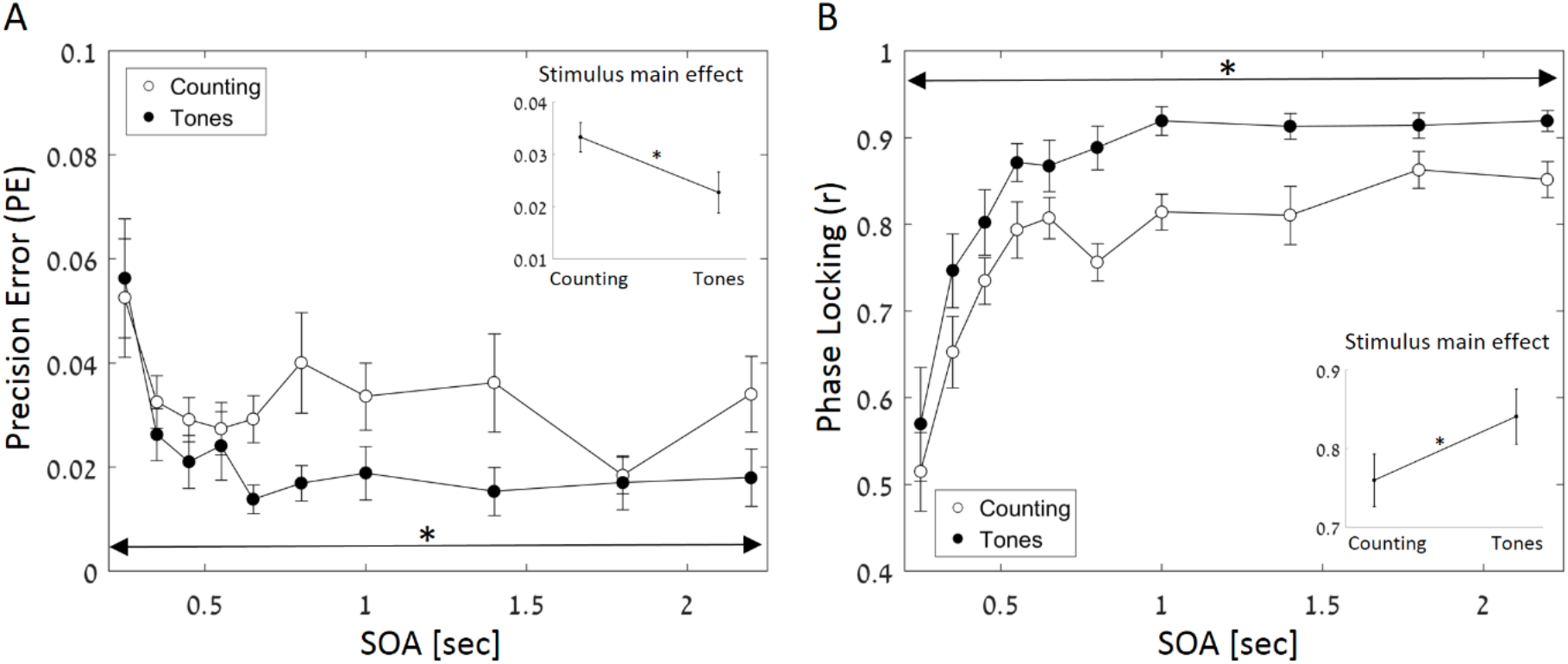
Synchronization tapping results. (A) Precision error and (B) Tapping Isochrony (phase-locking resultant), as a function of rate and stimulus type. Insets: Main effect of Stimulus Type on precision error and phase-locking. Error bars depict SEM.

Figure 4 shows the distribution of tapping times relative to each tone / digit in a sequence with an SOA of 0.8 sec, across all participants. For the tone stimuli (Figure 4A), tap times consistently precede the sound itself, which is consistent with previous studies (negative mean asynchrony; NMA) (Repp 1982; Repp and Su 2013). For the counting stimuli (Figure 4B), tap times were considerably more variable across digits, as participants tended to center their tap around the stressed syllable (which could vary from digit to digit) and not necessarily to the digit onset. This variability is characteristic of speech, which is naturally non-isochronous, and likely drives the reduced precision and isochrony when synchronizing to counting vs. tone sequences.

**Figure 4.**
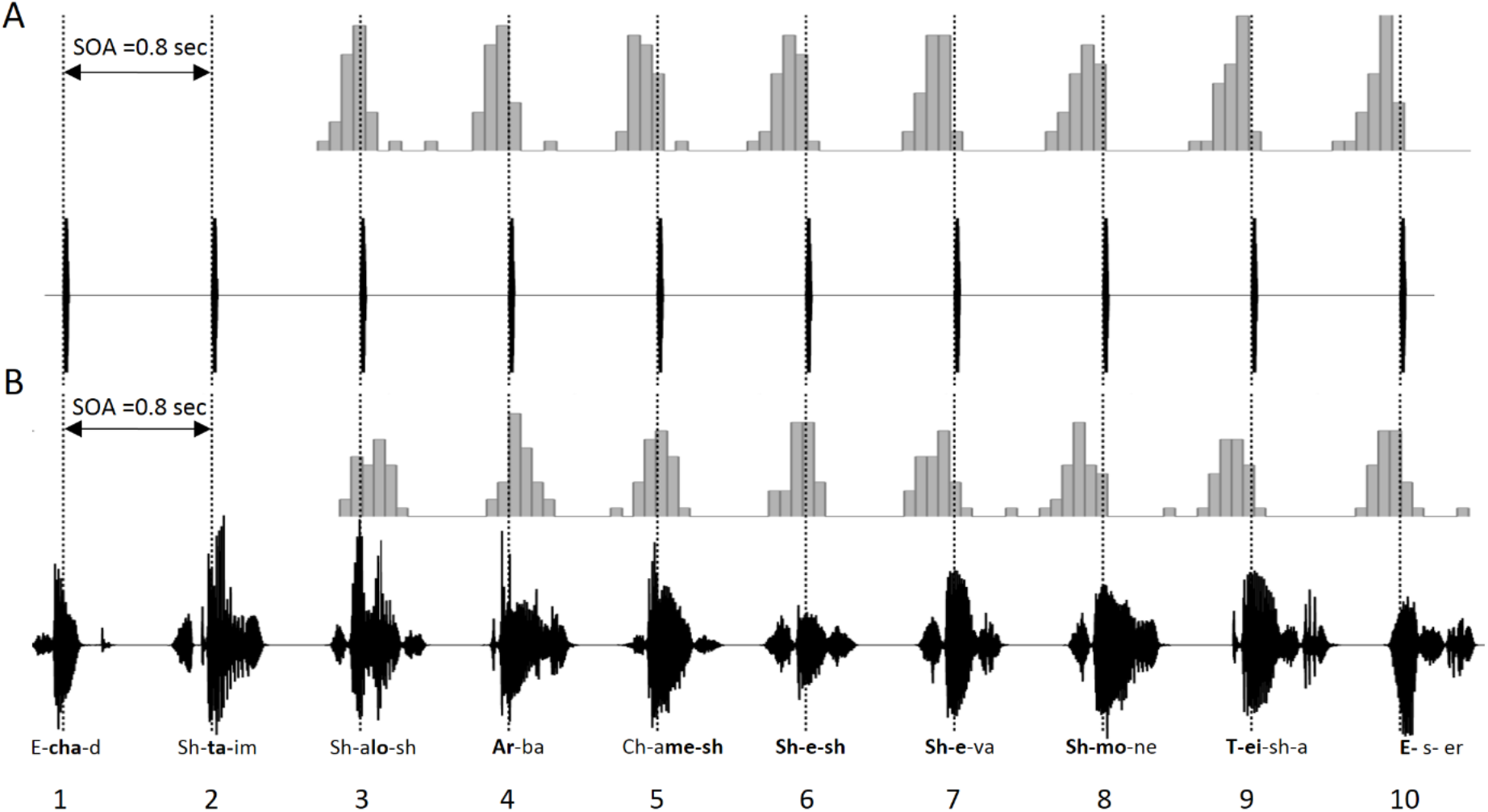
Distribution of tap-times around individual stimuli in the (A) tone and (B) counting sequence (example shown for SOA = 0.8 sec, which was the original counting recording). Dashed lines indicate the start time of each tone (panel A) or the times of metronome used to synchronize speech when recording the counting stimuli (panel B). The transliteration of each Hebrew digit is given at the bottom of panel B, with indication of the stressed syllable in bold.

### 3.2 Memory-paced Continuation-Tapping

For memory-paced continuation tapping, both precision error and isochrony showed a U-shape or inverse U-shape across rates, indicating that there is optimal performance for rates with SOAs of 0.55 and 0.65 sec [Figure 5, upper panel; main effect of rate: Precision Error: F(9,144)=10.1, p<0.001; phase-locking: F(9,144)=9.75, p<0.001]. Figure 6 shows a representative example of continuation tapping at three different rates from one participant. When comparing memory-paced tapping after passive listening vs. after active synchronization, we found a main effect of Task on both precision error [F(1,16)=11.5, p=0.004] and isochrony [F(1,16)=18.08, p<0.001] (Figure 5, middle panel), indicating better performance after active synchronization. This effect did not interact significantly with Rate [precision error: F(9,144)=1.86, p=0.061; isochrony: F(9,144)=0.73, p=0.67] or with Stimulus Type [precision error: F(1,16)=1.2, p=0.27; isochrony: F(1,16)=0.85, p=0.36], indicating that the profile of default rhythmic preferences was maintained for both passive listening and active synchronization.

**Figure 5.**
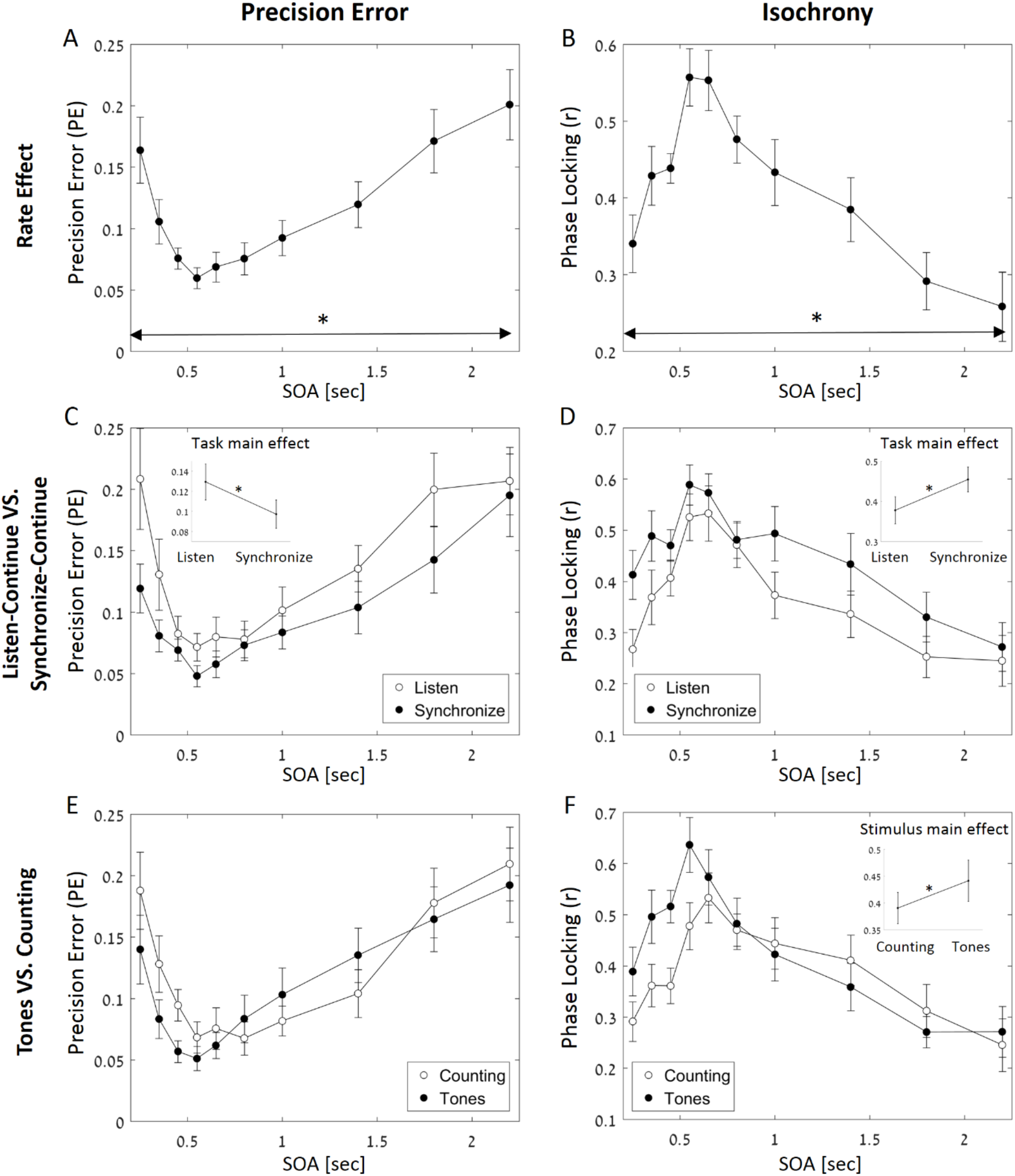
Continuation tapping results. <u>A&B.</u> Precision Error and Isochrony (phase-locking resultant) across all rates. Both measures show optimal performance for rates near 0.5 sec SOA. <u>C&D.</u> Precision Error and Isochrony across rates, separately for the Listen-Continue and Synchronize-Continue tasks. Both measures showed improved performance after active synchronization. <u>E&F.</u> Precision Error and Isochrony across rates, separately for the Tones and Counting stimuli. Isochrony was improved for tones, but similar precision error was observed for the two stimulus types. Insets: Main effects of Task on precision error and phase-locking (C&D) and of Stimulus Type on phase-locking (F). Error bars depict SEM

**Figure 6.**
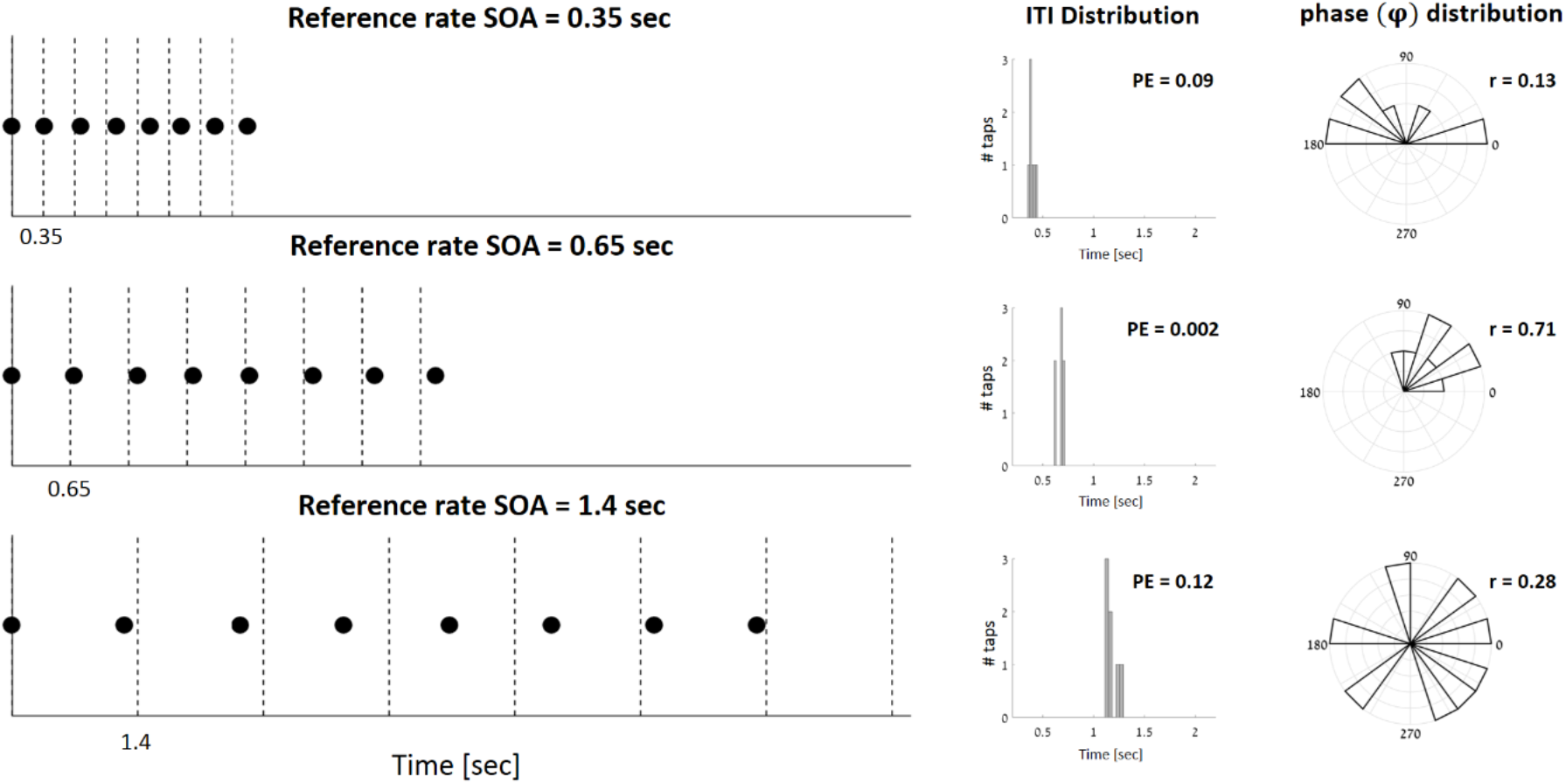
Example of Continuation tapping performance at three different rates: SOA = 0.35, 0.65 and 1.4 sec. Left: circles indicate tap times, with respect to the reference rhythm (dashed lines). Middle: ITI distribution and Precision Error (PE) for continuation tapping at each rate. Right: Phase distribution and resultant (r) value, reflecting the isochrony of continuation tapping at each rate.

Last, when comparing memory-paced tapping after hearing a rhythm that consisted of tones vs. counting, we found similar U-shape / inverted U-shape for precision error/phase locking across rates (respectively), for both Stimulus Types indicating a relatively similar range of optimal performance for rates around SOAs of ^~^0.45 - 0.65 sec (Figure 5, lower panel). We did find a significant main effect of Stimulus Type (Tones/Counting) on phase-locking [Figure 5F; F(1,16)=6.3, p=0.23], demonstrating more isochronous tapping after hearing a rhythm consisting of tones vs. counting. The main effect of Stimulus Type was not significant for precision error [Figure 5E; F(1,16)=2.09, p=0.167], but a significant interaction was found between Stimulus Type and Rate [F(9,144)=2.55, p=0.009] suggesting a shift in rhythmic preference for counting stimuli towards slightly slower rates (optimal range for tones: 0.45-0.65 sec SOA; optimal range for counting: 0.55-1 sec SOA).

## 4. Discussion

Here we studied the contribution of three factors to the human ability to synchronize with and reproduce rhythms in the environment: The role of motor engagement, the type of stimuli the rhythmic input is comprised of, and how performance varies across a broad range of rates. Consistent with previous findings, individuals displayed a large dynamic range for synchronization with near-perfect performance across all rates, with the exception of the extremely fast rhythms (SOA < 0.35 sec) (Kliger Amrani and Zion Golumbic 2020a). Similar synchronization accuracy was found for both simple tone rhythms and counting rhythms, which is indicative of the extraordinary flexibility of action-perception loops to adapt dynamically to the diverse range of rhythms in our environment (Repp and Steinman 2010; Chauvigné et al. 2014; Pérez-González and Malmierca 2014; Zamm et al. 2015). However, memory-paced continuation tapping, that relies on an internal representation of a previously-heard reference rhythm, is not as adaptable. The current results replicate the U-shape pattern reported previously, indicating a ‘sweet spot’ for optimal memory-paced tapping for rhythms with SOAs ^~^0.55 - 0.65 sec, and reduced performance for both faster and slower rates (Kliger Amrani and Zion Golumbic 2020a; Zalta et al. 2020). Interestingly, we found that this U-shape pattern is preserved for both tone and counting stimuli, and both when it was performed after active synchronization or passive listening. This pattern suggests that the internal representation of rhythms, and therefore also memory-paced tapping, is highly affected by default rhythm preferences. We also found that memory-paced tapping was significantly better after actively synchronizing to a reference rhythm vs. after passive listening. These results emphasize the importance of action-perception interactions in forming a stable temporal representation for rhythm. Last, synchronization and memory-paced continuation tapping performance were qualitatively similar for speech-derived rhythms and for simple tone sequences, though tapping to tones was generally more isochronous. This pattern broadens the conversation regarding the generalizability of the rhythmic entrainment hypothesis to more ecological rhythmic stimuli. We now turn to discuss the effects of motor-engagement, rhythmic defaults and stimulus type more in depth.

### 4.1 Importance of motor-engagement in memory-paced tapping

Scientists have long tried to model and understand the ‘internal clock inside our heads’, which enables us to reproduce time-intervals and rhythms from mental representations of past stimuli (Church 1984). Wing and Kristofferson proposed a model for memory-paced tapping pointing to combined contributions of perceptual and motor-based working memory (Wing and Kristofferson 1973). The importance of motor-engagement for rhythm perception has been substantiated by a wealth of neuroimaging studies, demonstrating the involvement of motor related regions – both cortical and subcortical – in timing- and rhythm-related processes, even in the absence of overt motion (Ivry 1996; Penhune et al. 1998; Lustig et al. 2005; Grahn and Brett 2007; Hassanpoor et al. 2012; Marvel et al. 2020). There are also growing indications that motor movement can directly affect auditory perception, particularly when stimuli are rhythmic or temporally predictable (Thaut et al. 2001; Schroeder et al. 2010; Chemin et al. 2014; Park et al. 2015; Morillon and Baillet 2017; Rimmele et al. 2018). Some examples for this are enhancement of perceptual thresholds (Reznik et al. 2021), increased sensitivity to auditory pitch patterns (Lange 2009; Morillon et al. 2014; Bauer et al. 2015) and enhanced neural representations for rhythmic sounds following movement (Chemin et al. 2014; Zalta et al. 2020). The current finding that memory-paced tapping is significantly better after actively synchronizing to a reference rhythm vs. after hearing it passively provides yet another indication for the strong influence of motor action on auditory rhythm perception. Specifically, it implies that not only does tapping *together with sounds* enhance perception of these sounds, but that motor engagement results in the formation of a more stable temporal representation in working memory, relative to the representation formed based on auditory input alone (Barkley et al. 1997; Chauvigné et al. 2014; Mioni et al. 2019).

### 4.2 Memory-paced tapping and rhythmic defaults

Motor contributions to auditory perception are proposedly brought about thorough coupling of motor and auditory cortices, acting together as a closed-loop action-perception system. This connectivity between the motor and auditory systems is plastic and amenable to improvement through training (Baumann et al. 2007), which lends itself to the improvement of fine and gross movements by music-supported therapy in patients with movement disorders (e.g. Parkinson’s and stroke patients; Thaut et al. 2001; Rodriguez-Fornells et al. 2012). Recent neurophysiological studies point specifically to nested neural oscillations in the delta (1-3Hz) and beta (15-25Hz) bands as the primary means for communication and transmission of efference copies between auditory and motor cortices (Stefanics et al. 2010; Arnal et al. 2015; Morillon and Baillet 2017; Morillon et al. 2019; Abbasi and Gross 2020). In particular, it is suggested that neural oscillations in auditory and motor cortices entrain to the rhythm of auditory stimuli, and thus become synchronized among themselves. Although in the current study we did not measure neural activity during the synchronization-continuation tasks, our results nonetheless carry some important insights regarding the underlying mechanisms generating the observed rhythmic behaviours. Specifically, they emphasize the existence and role of rhythmic defaults in the internal representation of rhythms and suggest a potential distinction in the operations of neural entrainment in synchronization vs. memory-paced tapping. As we observed, synchronization abilities were near-perfect for a vast range of rhythms – from ultra fast to ultra slow. A similarly large dynamic range for accurate synchronization has been observed for both simple and complex rhythms, even in young children, and is a testament to the flexibility of the action-perception loop to adapt to a wide-range of rhythms (Fraisse 1982; Large et al. 2002; Keller and Repp 2005; Jacoby and Repp 2012; Repp and Moseley 2012; Repp and Su 2013; Fujii et al. 2014). In comparison, memory-paced tapping is optimal for a substantially narrower range of rates, roughly between 0.55-0.65 sec SOA / ^~^1.5-1.8 Hz, resembling a resonance-like phenomenon. Our results converge with those of another recent study who demonstrated optimal beat discrimination capabilities for rates around ^~^1.6 Hz, after both passive listening and active tapping (Zalta et al. 2020). This ‘sweet spot’ for memory-paced tapping and discrimination converges with the documented range of spontaneous finger-tapping rhythms and of auditory-perceptual preference, which have been attributed to a ‘default’ internal oscillator underlying both the production and perception of rhythms (Styns and Leman 2007; McAuley 2010; Michaelis et al. 2014; Provasi et al. 2014; Large and Gray 2015b; Schwartze and Kotz 2015; Zamm et al. 2015; Scheurich et al. 2018). The current results suggest that, in contrast to synchronization, memory-paced tapping makes use of these rhythmic defaults, and if the reference rhythm falls within this optimal range, the default oscillator can be engaged to perpetuate its internal representation so it can ultimately be reproduced accurately. Interestingly, this preference was preserved even when memory-paced tapping was performed after passive listening without active motor engagement, further supporting the hypothesized generalization of rhythmic defaults across auditory perception and motor production (Fraisse 1982; Collyer et al. 1994; McAuley et al. 2006; Chen et al. 2008; Michaelis et al. 2014; Schwartze and Kotz 2015; Rose et al. 2020). (but see Kliger-Amrani & Zion Golumbic 2020 a&b regarding the limitations of this correspondence). However, our results also suggest that the farther the rhythm is from these defaults, the more difficult it is for the system to create stable memory representations for them, which may severely limit the utility of memory-based entrainment in the absence of ongoing stimulation.

### 4.3 Synchronization and memory-paced tapping to rhythmic speech

One of the domains in which entrainment has been hypothesized to play a functionally important role is speech processing. Motivated by the pseudo-rhythmic nature of speech, by the idea that synchronization can help guide perceptual resources towards points in time when meaningful information is delivered (Large and Jones 1999; Drake et al. 2000), as well as by the strong link between rhythmic capabilities and language skills (Corriveau and Goswami 2009; Woodruff Carr et al. 2014; Park et al. 2015), the possibility of neural entrainment to speech-rhythms has gained much enthusiasm in recent years (Cummins 2009; Peelle et al. 2012; Zion Golumbic et al. 2012; Ding and Simon 2014; Park et al. 2015; Keitel et al. 2018; Poeppel and Assaneo 2020). At the same time, there is also substantial skepticism on the matter, since although languages are often classified as “syllable - timed” or “stress - timed”, reflecting the idea that syllables/stressed-syllables occur at relatively stable time-intervals, natural speech is clearly not strictly isochronous (Dauer 1983; Lidji et al. 2011; Cummins 2015; Aubanel and Schwartz 2020). Moreover, it is not even clear which (if any) acoustic and/articulatory features drive the perception of speech as rhythmic (otherwise known as perceptual-center; or P-centers; Morton et al. 1976; De Jong 1994; Villing et al. 2011). This makes it difficult to test to what degree the notion of synchronization to rhythms in the environment extends to speech-derived rhythms (Haegens and Zion Golumbic 2018; Doelling and Assaneo 2021).

Here we take a step in that direction by using a speech stimulus that ***is*** (relatively) isochronous by nature: counting from 1 to 10. The adaptation of the synchronization-continuation task to counting rhythms allowed us to test whether individuals extrapolate the perception of an isochronous rhythm even when the sensory-elements constructing the sequence consist of speech-elements that vary in acoustic makeup, duration, stress location and other features that are characteristic of the acoustic variability of speech. Two previous studies that examined motor-tapping to speech rhythms found that individuals naturally synchronize their tapping to the P-centers in speech, at least in the case of *monosyllabic words* (Lidji et al. 2011; Villing et al. 2011). Another study that examined continuation motor tapping to tones while listening to French and German sentences, found that verbal changes were detected more efficiently when taps were congruently aligned to speech accents (P-centers), as compared to the incongruently aligned (Falk et al. 2017). Hence, although tapping to speech is a bit contrived, it allows us insight into the internal rhythmic representations that are derived from speech stimuli.

Not surprisingly, we found that both synchronization and memory-paced tapping are more isochronous and more precise for simple isochronous tone sequences relative to rhythmic counting. This is likely driven by the less strictly-rhythmic nature of speech rhythms, and the variability in duration and stress location across digits (Figure 4). At the same time, the effects of rate and of motor-engagement were qualitatively similar for the two types of stimuli. Specifically, for synchronization tapping participants showed high flexibility for adapting to a broad range of counting rhythms, and memory-paced tapping for counting followed a similar U-shaped pattern as was observed for tone sequences (albeit the range of ‘optimal’ counting rates included was slightly slower). This overlap in range of ‘optimal’ rates for tones and counting is in line with the proposition that rhythmic defaults underlying the representation of speech-rhythms and simple rhythms stem from a common oscillator (Assaneo et al. 2019, 2021). These results also converge with previous studies showing that there is an optimal range of rates where speech-intelligibility is optimal, inviting additional research into the potential commonalities between rhythm perception and speech processing (Lim 2010; Ghitza 2011; Peelle et al. 2012; Przybylski et al. 2013; Henry and Herrmann 2014; Doelling 2015; Adam Tierney, Travis White-Schwoch, Jessica MacLean 2017; Falk et al. 2017).

## 5. Conclusion

This study highlights the versatility and some of the constraints for entraining motor actions to rhythms in the environments. Despite the impressive flexibility that individuals display for synchronizing to a broad range of rhythms, it seems that maintaining these rhythms in working memory depends on internal rhythmic defaults, and as such is of a more limited nature. We also show that entrainment is possible to base on perceptual input only, but that temporal accuracy is substantially improved when the motor system is also engaged. Finally, this study contributes to the ongoing debates regarding the plausibility of entrainment to rhythms in speech, and shows that (at least for simple counting rhythms) performance is qualitatively similar to, but less accurate than, for simple tones sequences. Taken together, this work broadens the conversation about the factors affecting the way people interact with rhythms in their environment, and puts forth several specific and testable hypotheses regarding the underlying neural mechanisms.

## References

Abbasi O, Gross J. 2020. Beta-band oscillations play an essential role in motor–auditory interactions. Hum Brain Mapp. 41:656–665.

Adam Tierney, Travis White-Schwoch, Jessica MacLean and NKA. 2017. Individual Differences in Rhythm Skills: Links with Neural Consistency and Linguistic Ability. J Cogn Neurosci. 139.

Arnal LH, Doelling KB, Poeppel D. 2015. Delta-beta coupled oscillations underlie temporal prediction accuracy. Cereb Cortex. 25:3077–3085.

Assaneo MF, Rimmele JM, Sanz Perl Y, Poeppel D. 2021. Speaking rhythmically can shape hearing. Nat Hum Behav. 5:71–82.

Assaneo MF, Ripollés P, Orpella J, Lin WM, de Diego-Balaguer R, Poeppel D. 2019. Spontaneous synchronization to speech reveals neural mechanisms facilitating language learning. Nat Neurosci. 22:627–632.

Aubanel V, Schwartz JL. 2020. The role of isochrony in speech perception in noise. Sci Rep. 10:1–12.

Barkley R a, Koplowitz S, Anderson T, McMurray MB. 1997. Sense of time in children with ADHD: effects of duration, distraction, and stimulant medication. J Int Neuropsychol Soc. 3:359–369.

Bartlett NR, Bartlett SC. 1959. Synchronization of a motor response with an anticipated sensory event. Psychol Rev. 66:203–218.

Bauer AKR, Jaeger M, Thorne JD, Bendixen A, Debener S. 2015. The auditory dynamic attending theory revisited: A closer look at the pitch comparison task. Brain Res. 1626:198–210.

Baumann S, Koeneke S, Schmidt CF, Meyer M, Lutz K, Jancke L. 2007. A network for audio-motor coordination in skilled pianists and non-musicians. Brain Res. 1161:65–78.

Bravi R, Cohen EJ, Martinelli A, Gottard A, Minciacchi D. 2017. When non-dominant is better than dominant: Kinesiotape modulates asymmetries in timed performance during a synchronization-continuation task. Front Integr Neurosci. 11:1–16.

Breska A, Deouell LY. 2017. Neural mechanisms of rhythm-based temporal prediction: Delta phase-locking reflects temporal predictability but not rhythmic entrainment. PLOS Biol. 15:e2001665.

Chauvigné LAS, Gitau KM, Brown S. 2014. The neural basis of audiomotor entrainment: an ALE meta-analysis. Front Hum Neurosci. 8:776.

Chemin B, Mouraux A, Nozaradan S. 2014. Body Movement Selectively Shapes the Neural Representation of Musical Rhythms. Psychol Sci. 25:2147–2159.

Chen JL, Penhune VB, Zatorre RJ. 2008. Moving on time: brain network for auditory-motor synchronization is modulated by rhythm complexity and musical training. J Cogn Neurosci. 20:226–239.

Collyer CE, Broadbent HA, Church RM. 1994. Preferred rates of repetitive tapping and categorical time production. Percept Psychophys. 55:443–453.

Corriveau KH, Goswami U. 2009. Rhythmic motor entrainment in children with speech and language impairments: Tapping to the beat. Cortex. 45:119–130.

Cortese MD, Riganello F, Arcuri F, Pignataro LM, Buglione L. 2015. Rehabilitation of aphasia: Application of melodic-rhythmic therapy to Italian language. Front Hum Neurosci. 9:1–8.

Cravo AM, Rohenkohl G, Wyart V, Nobre AC. 2013. Temporal expectation enhances contrast sensitivity by phase entrainment of low-frequency oscillations in visual cortex. J Neurosci. 33:4002–4010.

Cummins F. 2009. Rhythm as entrainment: The case of synchronous speech. J Phon. 37:16–28.

Cummins F. 2015. Rhythm and Speech. Handb Speech Prod. 158–177.

Damm L, Varoqui D, De Cock VC, Dalla Bella S, Bardy B. 2020. Why do we move to the beat? A multi-scale approach, from physical principles to brain dynamics. Neurosci Biobehav Rev. 112:553–584.

Dauer RM. 1983. Stress-timing and syllable-timing reanalyzed. J Phon. 11:51–62.

De Jong KJ. 1994. The correlation of P-center adjustments with articulatory and acoustic events. Percept Psychophys. 56:447–460.

Ding N. 2016. Cortical Tracking of Hierarchical Linguistic Structures in Connected Speech. Nat Neurosci. 176:139–148.

Ding N, Simon JZ. 2014. Cortical entrainment to continuous speech: functional roles and interpretations. Front Hum Neurosci. 8:311.

Doelling. 2015. Acoustic landmarks drive delta-theta oscillations to enable speech comprehension by facilitating perceptual parsing. 85.

Doelling KB, Assaneo MF. 2021. Neural oscillations are a start toward understanding brain activity rather than the end. PLOS Biol. 19:e3001234.

Donnet S, Bartolo R, Fernandes JM, Cunha JPS, Prado L, Merchant H. 2014. Monkeys time their pauses of movement and not their movement-kinematics during a synchronization-continuation rhythmic task. J Neurophysiol. 111:2138–2149.

Drake C, Jones MR, Baruch C. 2000. The development of rhythmic attending in auditory sequences: Attunement, referent period, focal attending, Cognition.

Dunlap K. 1910. Reaction to rhythmic stimuli with attempt to synchronize. Psychol Rev. 17:399–416.

Falk S, Volpi-Moncorger C, Dalla Bella S. 2017. Auditory-motor rhythms and speech processing in French and German listeners. Front Psychol. 8:1–14.

Flach R. 2005. The transition from synchronization to continuation tapping. Hum Mov Sci. 24:465–483.

Fraisse P. 1982. Rhythm and Tempo. In: Deutsch D, editor. The Psychology of music. Academic Press. p. 149.

Fujii S, Watanabe H, Oohashi H, Hirashima M, Nozaki D, Taga G. 2014. Precursors of Dancing and Singing to Music in Three-to Four-Months-Old Infants. PLoS One. 9:e97680.

Gale DJ, Areshenkoff CN, Honda C, Johnsrude IS, Flanagan JR, Gallivan JP. 2021. Motor Planning Modulates Neural Activity Patterns in Early Human Auditory Cortex. Cereb Cortex. 31:2952–2967.

Ghitza O. 2011. Linking speech perception and neurophysiology: speech decoding guided by cascaded oscillators locked to the input rhythm. Front Psychol Audit Cogn Neurosci. 2:130:doi: 10.3389/fpsyg.2011.00130.

Grahn JA, Brett M. 2007. Rhythm and beat perception in motor areas of the brain. J Cogn Neurosci. 19:893–906.

Haegens S, Zion Golumbic EM. 2018. Rhythmic facilitation of sensory processing: A critical review. Neurosci Biobehav Rev. 86.

Harrison EC, Horin AP, Earhart GM. 2018. Internal cueing improves gait more than external cueing in healthy adults and people with Parkinson disease. Sci Rep. 8:15525.

Hassanpoor H, Fallah A, Raza M. 2012. New role for astroglia in learning: Formation of muscle memory. Med Hypotheses. 79:770–773.

Henry MJ, Herrmann B. 2014. Low-Frequency Neural Oscillations Support Dynamic Attending in Temporal Context. Timing Time Percept. 2:62–86.

Hove MJ, Gravel N, Spencer RMC, Valera EM. 2017. Finger tapping and pre-attentive sensorimotor timing in adults with ADHD. Exp Brain Res. 235:3663–3672.

Ivry RB. 1996. The representation of temporal information in perception and motor control. Curr Opin Neurobiol. 6:851–857.

Ivry RB, Richardson TC. 2002. Temporal control and coordination: The multiple timer model. Brain Cogn. 48:117–132.

Jacoby N, Repp BH. 2012. A general linear framework for the comparison and evaluation of models of sensorimotor synchronization. Biol Cybern. 106:135–154.

Jaeger M, Bleichner MG, Bauer A-KR, Mirkovic B, Debener S. 2018. Did You Listen to the Beat? Auditory Steady-State Responses in the Human Electroencephalogram at 4 and 7 Hz Modulation Rates Reflect Selective Attention. Brain Topogr.

Jones MR. 2019. Time will tell: A theory of dynamic attending. Oxford University Press.

Keitel A, Gross J, Kayser C. 2018. Perceptually relevant speech tracking in auditory and motor cortex reflects distinct linguistic features. PLOS Biol. 16:e2004473.

Keller PE, Repp BH. 2005. Staying offbeat: Sensorimotor syncopation with structured and unstructured auditory sequences. Psychol Res Psychol Forsch. 69:292–309.

Kliger Amrani A, Zion Golumbic E. 2020a. Spontaneous and stimulus-driven rhythmic behaviors in ADHD adults and controls. Neuropsychologia. 146.

Kliger Amrani A, Zion Golumbic E. 2020b. Testing the stability of ‘Default’ motor and auditory-perceptual rhythms–A replication failure dataset. Data Br. 32:106044.

Kornysheva K, Schubotz RI. 2011. Impairment of Auditory-Motor Timing and Compensatory Reorganization after Ventral Premotor Cortex Stimulation. PLoS One. 6:e21421.

Kösem A, Bosker HR, Takashima A, Meyer A, Jensen O, Hagoort P. 2018. Neural Entrainment Determines the Words We Hear. Curr Biol. 28:2867–2875.e3.

Koshimori Y, Strafella AP, Valli M, Sharma V, Cho S, Houle S, Thaut MH. 2019. Motor Synchronization to Rhythmic Auditory Stimulation (RAS) Attenuates Dopaminergic Responses in Ventral Striatum in Young Healthy Adults: [11C]-(+)-PHNO PET Study. Front Neurosci. 13:106.

Lakatos P, Karmos G, Mehta AD, Ulbert I, Schroeder CE. 2008. Entrainment of Neuronal Oscillations as a Mechanism of Attentional Selection. Science (80-). 320:110–113.

Lakatos P, Musacchia G, O’Connel MN, Falchier AY, Javitt DC, Schroeder CE. 2013. The spectrotemporal filter mechanism of auditory selective attention. Neuron. 77:750–761.

Lakatos P, Shah AS, Knuth KH, Ulbert I, Karmos G, Schroeder CE. 2005. An Oscillatory Hierarchy Controlling Neuronal Excitability and Stimulus Processing in the Auditory Cortex. J Neurophysiol. 94:1904–1911.

Lange K. 2009. Brain correlates of early auditory processing are attenuated by expectations for time and pitch. Brain Cogn. 69:127–137.

Large EW. 2008. Resonating to musical rhythm: theory and experiment, Psychology of time.

Large EW, Fink P, Kelso JAS. 2002. Tracking simple and complex sequences. Psychol Res. 66:3–17.

Large EW, Gray PM. 2015a. Spontaneous Tempo and Rhythmic Entrainment in a Bonobo (Pan paniscus). 129:317–328.

Large EW, Gray PM. 2015b. Spontaneous tempo and rhythmic entrainment in a bonobo (Pan paniscus). J Comp Psychol. 129:317–328.

Large EW, Jones MR. 1999. The dynamics of attending: How people track time-varying events. Psychol Rev. 106:119–159.

Lewis PA, Wing AM, Pope PA, Praamstra P, Miall RC. 2004. Brain activity correlates differentially with increasing temporal complexity of rhythms during initialisation, synchronisation, and continuation phases of paced finger tapping. Neuropsychologia. 42:1301–1312.

Lidji P, Palmer C, Peretz I, Morningstar M. 2011. Listeners feel the beat: Entrainment to English and French speech rhythms. Psychon Bull Rev. 18:1035–1041.

Lim HA. 2010. Effect of “developmental speech and language training through music” on speech production in children with autism spectrum disorders. J Music Ther. 47:2–26.

Loukina A, Kochanski G, Rosner B, Keane E, Shih C. 2011. Rhythm measures and dimensions of durational variation in speech. J Acoust Soc Am. 129:3258–3270.

Lustig C, Matell MS, Meck WH. 2005. Not “just” a coincidence: Frontal-striatal interactions in working memory and interval timing. Memory. 13:441–448.

Marvel CL, Morgan OE, Kronemer SI, Haven N, Program N, Haven N. 2020. How the Motor System Integrates with Working Memory. Neurosci Biobehav. 184–194.

Marvel CL, Morgan OP, Kronemer SI. 2019. How the motor system integrates with working memory. Neurosci Biobehav Rev. 102:184–194.

McAuley JD. 2010. Tempo and Rhythm. Springer, New York, NY. p. 165–199.

McAuley JD, Jones MR, Holub S, Johnston HM, Miller NS. 2006. The time of our lives: Life span development of timing and event tracking. J Exp Psychol Gen. 135:348–367.

McNab F, Klingberg T. 2008. Prefrontal cortex and basal ganglia control access to working memory. Nat Neurosci. 11:103–107.

Merchant H, Zarco W, Pérez O, Prado L, Bartolo R. 2011. Measuring time with different neural chronometers during a synchronization-continuation task. Proc Natl Acad Sci U S A. 108:19784–19789.

Michaelis K, Wiener M, Thompson JC. 2014. Passive listening to preferred motor tempo modulates corticospinal excitability. Front Hum Neurosci. 8:1–10.

Mioni G, Capodieci A, Biffi V, Porcelli F, Cornoldi C. 2019. Difficulties of children with symptoms of attention-deficit/hyperactivity disorder in processing temporal information concerning everyday life events. J Exp Child Psychol. 182:86–101.

Morillon B, Arnal LH, Schroeder CE, Keitel A. 2019. Prominence of delta oscillatory rhythms in the motor cortex and their relevance for auditory and speech perception. Neurosci Biobehav Rev. 107:136–142.

Morillon B, Baillet S. 2017. Motor origin of temporal predictions in auditory attention. Proc Natl Acad Sci U S A. 114:E8913–E8921.

Morillon B, Schroeder CE, Wyart V. 2014. Motor contributions to the temporal precision of auditory attention. Nat Commun. 5.

Morton J, Marcus S, Frankish C. 1976. Perceptual centers (P-centers). Psychol Rev. 83:405–408.

Nenadic I, Gaser C, Volz H-P, Rammsayer T, Häger F, Sauer H. 2003. Processing of temporal information and the basal ganglia: new evidence from fMRI. Exp Brain Res. 148:238–246.

Nozaradan S, Peretz I, Keller PE. 2016. Individual Differences in Rhythmic Cortical Entrainment Correlate with Predictive Behavior in Sensorimotor Synchronization. Sci Rep. 6:20612.

Palmer C, Lidji P, Peretz I. 2014. Losing the beat: deficits in temporal coordination. Philos Trans R Soc B Biol Sci. 369:20130405.

Park H, Ince RAA, Schyns PG, Thut G, Gross J. 2015. Frontal Top-Down Signals Increase Coupling of Auditory Low-Frequency Oscillations to Continuous Speech in Human Listeners. Curr Biol. 25:1649–1653.

Peelle JE, Gross J, Davis MH. 2012. Phase-Locked Responses to Speech in Human Auditory Cortex are Enhanced During Comprehension. Cereb Cortex.

Penhune VB, Zattore RJ, Evans a C. 1998. Cerebellar contributions to motor timing: a PET study of auditory and visual rhythm reproduction. J Cogn Neurosci. 10:752–765.

Pérez-González D, Malmierca MS. 2014. Adaptation in the auditory system: an overview. Front Integr Neurosci. 8:19.

Phillips-Silver J, Toiviainen P, Gosselin N, Piché O, Nozaradan S, Palmer C, Peretz I. 2011. Born to dance but beat deaf: A new form of congenital amusia. Neuropsychologia. 49:961–969.

Poeppel D, Assaneo MF. 2020. Speech rhythms and their neural foundations. Nat Rev Neurosci. 21:322–334.

Provasi J, Anderson DI, Barbu-Roth M. 2014. Rhythm perception, production, and synchronization during the perinatal period. Front Psychol. 5:1048.

Przybylski L, Bedoin N, Krifi-Papoz S, Herbillon V, Roch D, Léculier L, Kotz SA, Tillmann B. 2013. Rhythmic auditory stimulation influences syntactic processing in children with developmental language disorders. Neuropsychology. 27:121–131.

Repp B, Steinman S. 2010. Simultaneous event-based and emergent timing: Synchronization, continuation, and phase correction. J Mot Behav. 42:111–126.

Repp BH. 1982. Phonetic trading relations and context effects: new experimental evidence for a speech mode of perception. Psychol Bull. 92:81–110.

Repp BH, Moseley GP. 2012. Anticipatory phase correction in sensorimotor synchronization. Hum Mov Sci. 31:1118–1136.

Repp BH, Su Y-H. 2013. Sensorimotor synchronization: A review of recent research (2006–2012). Psychon Bull Rev. 20:403–452.

Reznik D, Hachohen N, Buaron B, Zion-Golumbic EM, Mukamel R. 2021. Motor-evoked neural responses in auditory cortex are associated with improved sensitivity to self-generated sounds. Cereb Cortex. in press.

Rimmele JM, Morillon B, Poeppel D, Arnal LH. 2018. Proactive Sensing of Periodic and Aperiodic Auditory Patterns. Trends Cogn Sci. 22:870–882.

Rodriguez-Fornells A, Rojo N, Amengual JL, Ripollés P, Altenmüller E, Münte TF. 2012. The involvement of audio-motor coupling in the music-supported therapy applied to stroke patients. Ann N Y Acad Sci. 1252:282–293.

Roerdink M, Bank PJM, Peper CLE, Beek PJ. 2011. Walking to the beat of different drums: Practical implications for the use of acoustic rhythms in gait rehabilitation. Gait Posture. 33:690–694.

Rohenkohl G, Cravo a. M, Wyart V, Nobre a. C. 2012. Temporal Expectation Improves the Quality of Sensory Information. J Neurosci. 32:8424–8428.

Rose D, Cameron DJ, Lovatt PJ, Grahn JA, Annett LE. 2020. Comparison of Spontaneous Motor Tempo during Finger Tapping, Toe Tapping and Stepping on the Spot in People with and without Parkinson’s Disease. J Mov Disord. 13:47–56.

Rosen S. 1992. Temporal Information in Speech: Acoustic, Auditory and Linguistic Aspects. Phil Trans Biol Sci. 336:367–373.

Rosen S, Fourcin A. 1986. Frequency selectivity and the perception of speech. In: Frequency Selectivity in Hearing. London: Academic Press Inc. p. 373–397.

Scheurich R, Zamm A, Palmer C. 2018. Tapping Into Rate Flexibility: Musical Training Facilitates Synchronization Around Spontaneous Production Rates. Front Psychol. 9:458.

Schmidt R, Herrojo Ruiz M, Kilavik BE, Lundqvist M, Starr PA, Aron AR. 2019. Beta Oscillations in Working Memory, Executive Control of Movement and Thought, and Sensorimotor Function. J Neurosci. 39:8231–8238.

Schön D, Tillmann B. 2015. Short- and long-term rhythmic interventions: Perspectives for language rehabilitation. Ann N Y Acad Sci. 1337:32–39.

Schroeder CE, Lakatos P. 2009. Low-frequency neuronal oscillations as instruments of sensory selection. Trends Neurosci. 32:9–18.

Schroeder CE, Wilson DA, Radman T, Scharfman H, Lakatos P. 2010. Dynamics of Active Sensing and perceptual selection. Curr Opin Neurobiol. 20:172–176.

Schwartze M, Keller PE, Kotz SA. 2016. Spontaneous, synchronized, and corrective timing behavior in cerebellar lesion patients. Behav Brain Res. 312:285–293.

Schwartze M, Keller PE, Patel AD, Kotz SA. 2011. The impact of basal ganglia lesions on sensorimotor synchronization, spontaneous motor tempo, and the detection of tempo changes. Behav Brain Res. 216:685–691.

Schwartze M, Kotz SA. 2015. The Timing of Regular Sequences: Production, Perception, and Covariation. J Cogn Neurosci. 27:1697–1707.

Serrien DJ. 2008. The neural dynamics of timed motor tasks: Evidence from a synchronization-continuation paradigm. Eur J Neurosci. 27:1553–1560.

Sowiński J, Dalla Bella S. 2013. Poor synchronization to the beat may result from deficient auditory-motor mapping. Neuropsychologia. 51:1952–1963.

Stefanics G, Hangya B, HernÃ¡di I, Winkler I, Lakatos P, Ulbert I. 2010. Phase Entrainment of Human Delta Oscillations Can Mediate the Effects of Expectation on Reaction Speed. J Neurosci. 30:13578–13585.

Stevens LT. 1886. On The Time-Sense. Oxford Univ Press. 11:393–404.

Styns F, Leman M. 2007. Walking on music. Hum Mov Sci. 26:769–785.

Teki S, Gu B-M, Meck WH. 2017. The Persistence of Memory: How the Brain Encodes Time in Memory. Curr Opin Behav Sci. 17:178–185.

Thaut MH, McIntosh KW, McIntosh GC, Hoemberg V. 2001. Auditory rhythmicity enhances movement and speech motor control in patients with Parkinson’s disease. Funct Neurol. 16:163–172.

Tierney A, Kraus N. 2013. The ability to move to a beat is linked to the consistency of neural responses to sound. J Neurosci. 33:14981–14988.

Tierney A, Kraus N. 2014. Auditory-motor entrainment and phonological skills: precise auditory timing hypothesis (PATH). Front Hum Neurosci. 8:1–9.

Toyomura A, Shibata M, Kuriki S. 2012. Self-paced and externally triggered rhythmical lower limb movements: A functional MRI study. Neurosci Lett. 516:39–44.

Tranchant P, Vuvan DT, Peretz I. 2016. Keeping the Beat: A Large Sample Study of Bouncing and Clapping to Music. PLoS One. 11:e0160178.

Villing RC, Repp BH, Ward TE, Timoney JM. 2011. Measuring perceptual centers using the phase correction response. Attention, Perception, Psychophys. 73:1614–1629.

Wiener M, Turkeltaub P, Coslett HB. 2010. The image of time: A voxel-wise meta-analysis. Neuroimage. 49:1728–1740.

Wing AM, Kristofferson a. B. 1973. The timing of interresponse intervals. Percept Psychophys. 13:455–460.

Woodruff Carr K, White-Schwoch T, Tierney AT, Strait DL, Kraus N. 2014. Beat synchronization predicts neural speech encoding and reading readiness in preschoolers. Proc Natl Acad Sci U S A. 111:14559–14564.

Zalta A, Petkoski S, Morillon B. 2020. Natural rhythms of periodic temporal attention. Nat Commun. 11.

Zamm A, Pfordresher PQ, Palmer C. 2015. Temporal coordination in joint music performance: effects of endogenous rhythms and auditory feedback. Exp Brain Res. 233:607–615.

Zamm A, Wang Y, Palmer C. 2018. Musicians’ Natural Frequencies of Performance Display Optimal Temporal Stability. J Biol Rhythms. 33:432–440.

Zion Golumbic EM, Poeppel D, Schroeder CE. 2012. Temporal Context in Speech Processing and Attentional Stream Selection: A Behavioral and Neural perspective. Brain Lang. 122:151–161.

